# Viral population genomics reveals host and infectivity impact on SARS-CoV-2 adaptive landscape

**DOI:** 10.1101/2021.12.30.474516

**Authors:** Kaitlyn Gayvert, Richard Copin, Sheldon McKay, Ian Setliff, Wei Keat Lim, Alina Baum, Christos A. Kyratsous, Gurinder S. Atwal

**Affiliations:** Regeneron Pharmaceuticals, Inc., Tarrytown, NY 10591

## Abstract

Public health surveillance, drug treatment development, and optimization of immunological interventions all depend on understanding pathogen adaptation, which differ for specific pathogens. SARS-CoV-2 is an exceptionally successful human pathogen, yet complete understanding of the forces driving its evolution is lacking. Here, we leveraged almost four million SARS-CoV-2 sequences originating mostly from non-vaccinated naïve patients to investigate the impact of functional constraints and natural immune pressures on the sequence diversity of the SARS-CoV-2 genome. Overall, we showed that the SARS-CoV-2 genome is under strong and intensifying levels of purifying selection with a minority of sites under diversifying pressure. With a particular focus on the spike protein, we showed that sites under selection were critical for protein stability and virus fitness related to increased infectivity and/or reduced neutralization by convalescent sera. We investigated the genetic diversity of SARS-CoV-2 B and T cell epitopes and determined that the currently known T cell epitope sequences were highly conserved. Outside of the spike protein, we observed that mutations under selection in variants of concern can be associated to beneficial outcomes for the virus. Altogether, the results yielded a comprehensive map of all sites under selection across the entirety of SARS-CoV-2 genome, highlighting targets for future studies to better understand the virus spread, evolution and success.

## INTRODUCTION

The coronavirus disease 2019 (COVID-19) global pandemic has motivated a widespread effort to understand adaptation mechanisms of its etiologic agent, the severe acute respiratory syndrome coronavirus 2 (SARS-CoV-2). As a result, scientists and medical professionals from around the world have sequenced the SARS-CoV-2 genome from patient isolates and disseminated their findings at unprecedented speed through curated data repositories such as the global initiative on sharing all influenza data (GISAID. https://www.gisaid.org)^1,2^. This provided a unique dataset critical to determine transmission patterns and identify variants that may be associated with virulence and disease severity. This has become even more critical with the emergence of various variants under surveillance (VUS), which are associated with increased transmissibility and the potential to escape from natural immunity, therapeutic antibodies, and vaccine mediated immunity^3^.

To date, we still know little about the forces driving the conservation and diversity of SARS-COV-2 genome and the potential implication on treatment development and efficacy. While mutations emerge randomly in the genome, the accumulation or loss of variants can be beneficial to the virus within and across the host environment. This genetic adaptation can lead to both diversification or conservation of viral protein domains or amino acids and is key to the successful spread of SARS-CoV-2 in the human population. Selection driving amino acid conservation is called purifying or negative selection, and tends to conserve the integrity of a domain structure or function. Conversely, selection leading to the accumulation of amino acid changes is called positive selection and may be associated with gain of function through molecular alterations. During infection, pressure to escape recognition by antigen-targeting immune factors, such as antibodies and T lymphocytes, often leads to positive selection of viral antigens. The consequences of antigenic variation can include persistent viral infection^4^, pandemics of diseases^5^, and reinfection after recovery^6,7^. Antigenic variation can also impact therapeutics or vaccine efficacy, as it confounds development of treatments and vaccines that can protect against diverse viral strains ^7-9^. Thus, characterizing which regions in the genome are conserved or diverse with an emphasis on antigenic targets has critical implications for drug development and resistance.

To date, both, B cell and T cell immune responses have been reported to play roles in controlling SARS-CoV-2 infections^10^. B cells produce both anti-SARS-CoV-2 neutralizing and non-neutralizing antibodies, which comprise the majority of the polyclonal responses. Many of the most potently neutralizing anti-SARS-CoV-2 antibodies target the receptor-binding domain (RBD) of the viral spike protein, often competing with its binding to the ACE2 cell host receptor^11-21^. Antigen-specific T cells recognize short peptide epitopes generated by proteolysis of pathogen proteins, bound to HLA molecules on the surface of antigen-presenting cells. SARS-CoV-2 reactive memory CD4+^22^ and CD8+^23^ T cells have also been reported in unexposed individuals, suggesting that pre-existing cross-reactive T cell could drive disease outcomes in infected patients.

Understanding the forces shaping viral genomes in the past have helped better informed treatment development against life threatening infections. For example, evolutionary analyses of vector-borne RNA viruses, including dengue^24^, West Nile Virus^25^, and Zika^26^, have reported extensive levels of purifying selection, showing little impact of the immune system on antigenic variation and suggesting long-established adaptation to the natural host. By contrast, it was shown that the Ebola genome was under little selective pressure, indicating that the genome diversity was the unique result of genetic drift^27^. Finally, analyses of influenza sequences from the 2009 H1N1 pandemic reported that only the fraction of the genome harboring antigenic targets was under diversifying selection^28^.

To understand the forces shaping SARS-CoV-2 genome, and better inform treatment development, we analyzed an extensive dataset of 3,884,234 SARS-CoV-2 genome sequences available in GISAID. As the first vaccine against SARS-COV-2 was administered in December 2020 and distribution ramped up significantly only during spring 2021, the vast majority of these sequences pre-date the vaccine era. As such, they represent a great opportunity to identify and understand the impact of natural selection on sequence diversity of SARS-CoV-2 in naïve hosts. First, we established the evolutionary relationship of SARS-CoV-2 isolates and highlighted amino acid variants spreading in the community at high frequency. We then used evolutionary analysis to determine selection forces driving genetic diversity in SARS-CoV-2 at both protein and amino acid levels. Finally, we present a comprehensive genetic analysis of the currently known SARS-CoV-2 B and T cell epitopes to determine whether all SARS-CoV-2 antigens represent potential target of immune escape.

## RESULTS

### Expanded Analysis of Sequence Variation in SARS-CoV-2

Since the first SARS-CoV-2 genome sequence was reported in early January 2020, there have been almost four million sequences deposited to GISAID as of late September 2021 (https://www.gisaid.org/)^1,2^. Each genome sequence is associated with comprehensive patient-related metadata that can be used to determine the time of infection and the geographical origin of the virus isolates (Figure 1A). We compared the identity of all coding sequences retrieved from 3,884,234 curated genomes. We found that 41% of all variants were identified in only one given isolate (singleton). Since singletons may be transient and destined to be removed by purifying selection, their biological significance is uncertain. However, there were 1,689 amino acid changes shared across at least 2,000 isolates (high frequency variants or HFV), with a disproportionate number coming from the ORF7a (n=61; out of 121 possible sites), ORF8 (n=51/121), and ORF3a (n=107/275). Because the spike protein is the focus of most drug development efforts, characterization of its genetic diversity is critical for clinical surveillance and treatment outcome prediction. HFV identified within the spike protein were found accumulating within the N-terminal domain (NTD) (31%; 90 / 291) (Figure 1B). The RBD had a lower number of HFV (9%; 20 / 223) when compared to the full spike protein (16%; 204 / 1,273) or to the SARS-CoV-2 genome (15%; 1,456 / 9,701) as a whole. Overall, the spike protein did not show any enrichment of sequence diversity compared to the SARS-CoV-2 entire genome sequence.

**FIGURE 1.**
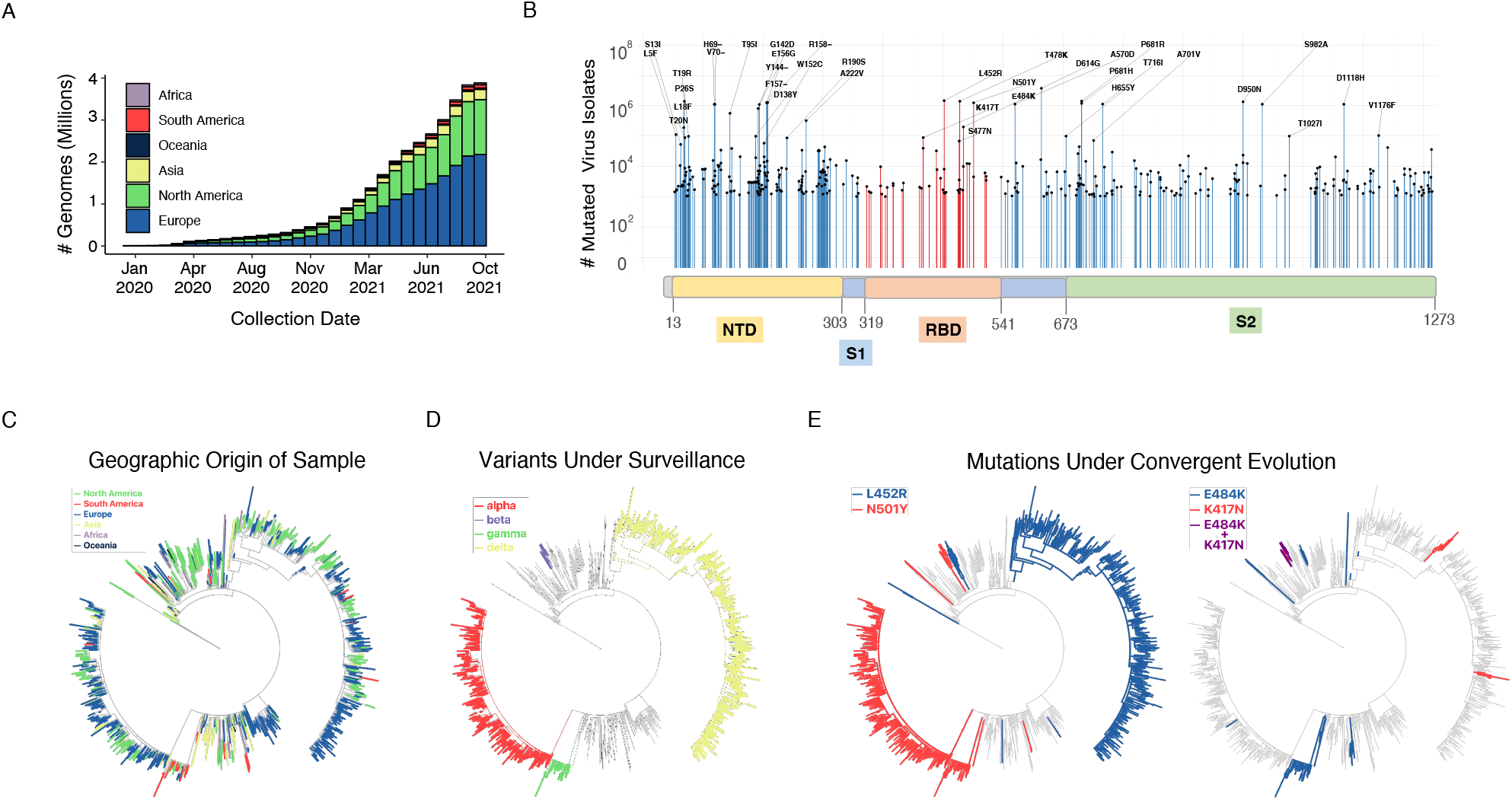
(A) Cumulative count of SARS-CoV-2 genome sequences in GISAID by collection date and continent of sample origin as of September 26, 2021. (B) Distribution of high frequency variants in the spike protein sequence. The RBD and NTD domains are highlighted orange and yellow, respectively. For a number of residues with high deviation from wild type sequence (417, 452, 484, 501, 614), the most common mutation is indicated. (C) SARS-CoV-2 maximum-likelihood phylogeny of high-diversity 1000-genome subset. The tree is outgroup rooted with the related bat CoV genome RATG13. The root branch length is truncated to emphasize relationships amongst SARS-CoV-2 genomes. The geographic origin of each of the sequences is denoted by branch color. (D) Distribution of haplotypes of the 4 variants of concern (alpha, beta, gamma, delta), mapped onto the phylogeny. (E) Distribution of mutations under convergent evolution (L452R, E484K), mapped onto the phylogeny.

### SARS-CoV-2 diversity across phylogenetic lineages

Detailed, rigorous analysis of a data set of four million genomes is challenging with current computing resources. To obtain computationally feasible data set, we developed a down-sampling method to select a subset of 20,000 genomes (see Methods)^29,30^. Genome-based phylogenetic analysis further resolved relationships among samples and the evolution of the virus in relation to geographic distribution. Many lineages contained SARS-CoV-2 isolates from several continents, indicating limited geographical clustering consistent with a global pandemic (Figure 1C). Comparison of the diversity of individual genomes by lineage allowed identification of mutations exhibiting several instances of convergent evolution (Figure 1D). These mutations included L452R and E484K, which were identified independently in genomes from at least 5 and 7 lineages, respectively. This indicates that emergence of both mutations occurred as a result of similar selective forces in independent individuals rather than linear transmission. Because phylogeny-defining mutations can be critical to understand virus biology, a further examination of positive and purifying selection and functional studies of how variants may impact SARS-CoV-2 biology was warranted.

### Selective landscape of the SARS-CoV-2 genome

Current Bayesian estimates of the SARS-CoV-2 genome mutation rate are approximately 7×10^−4^ nucleotide substitutions per site per year^31,32^, which is relatively low compared to other human coronaviruses and RNA viruses^32^. The low mutation rate may suggest that purifying selection dominates the SARS-CoV-2 genome. However, it does not rule out the possibility that some amino acid sites may be under positive selection. To characterize the selective forces driving SARS-CoV-2 gene sequence diversity, we applied a Bayesian codon model, FUBAR^30^, to identify sites evolving under natural selection. Codon substitution models estimate site-specific synonymous (dS, alpha) and nonsynonymous (dS, beta) substitution rates from a given sequence alignment and phylogeny, which in turn are used to identify sites that are evolving significantly faster (alpha<beta; positive selection) or slower (alpha>beta; purifying selection) than expected. Quickly spreading mutations under positive selection, especially those that arise independently across multiple lineages via convergent evolution, can be identified through these approaches by their elevated dN/dS rates (>>1). Similarly, sites where the reference alleles are conserved and mutations fall exclusively on shallow branches of the phylogeny are typically under purifying selection, as evidenced by low dN/dS ratios (<<1), and are more likely to negatively impact the fitness of the virus ^33^.

Overall, a gene-based measure of selection confirmed strong purifying selection acting on the SARS-CoV-2 genome but also identified a substantial number of sites under positive selection in several genes (Figure 2A). Amongst structural and accessory protein-encoding genes, *N, ORF3a, ORF7a, ORF8, and ORF10* genes showed an elevated number of sites under positive selection, indicating that these genes were under distinct selective forces when compared to the rest of the genome. Early stop codons are amongst the most frequent HFV in ORF3a (Q57H) and ORF8 (Q27*)^34^ and ORF10 is no longer treated as a protein-coding gene^35^. Interestingly, most of these genes have been implicated in the suppression of the innate immune response, particularly in the suppression of type one interferon signaling (ORF7a^36^, ORF8), blocking autophagy (ORF3a^37^, ORF7a^36^), and downregulation of antigen presentation (ORF8^38^).

**FIGURE 2.**
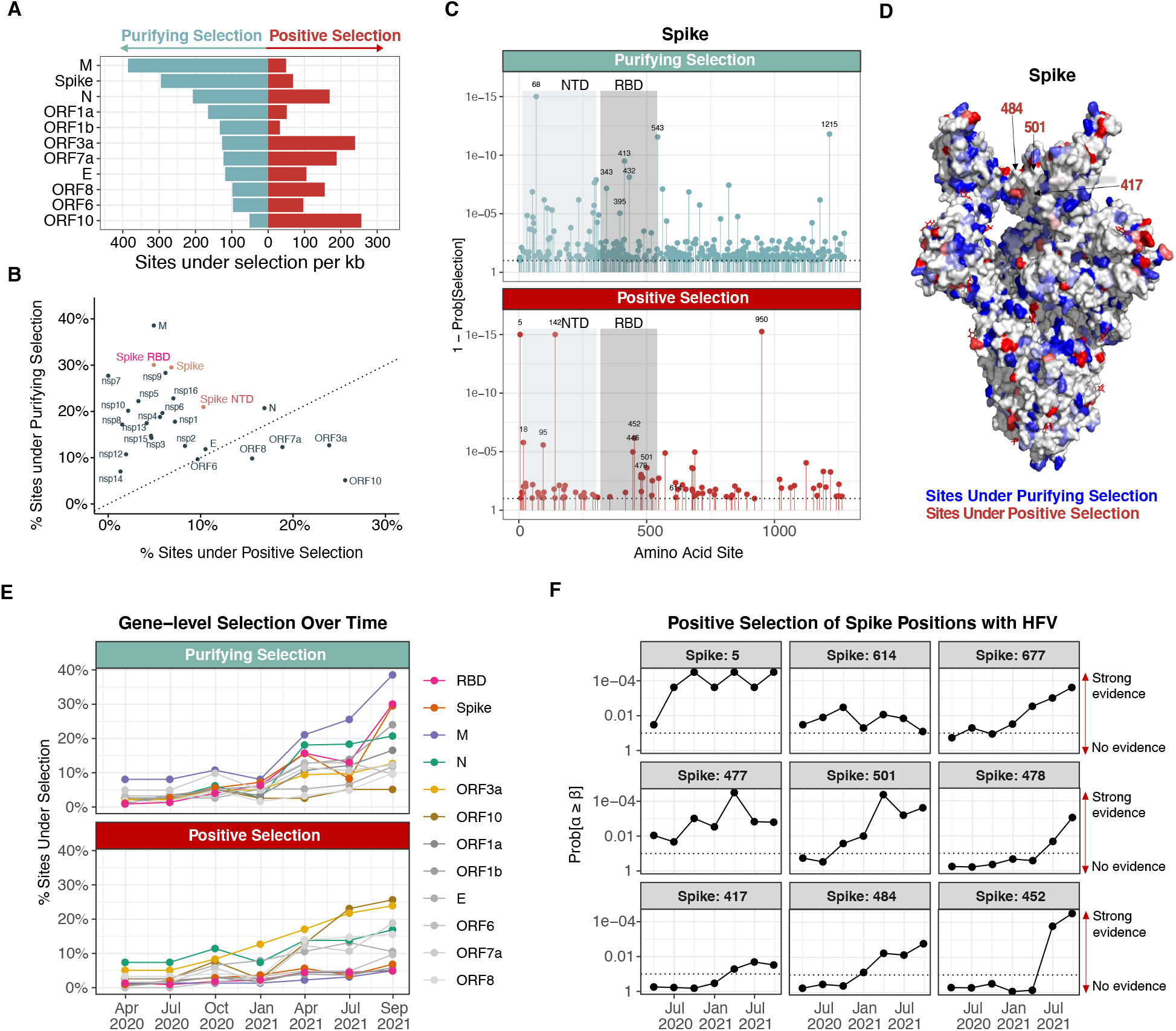
Amino acid sites under natural selection in SARS-CoV-2 genome. (A) Gene length normalized counts (counts per kb) of sites under significant purifying selection (light blue bars) and positive selection (red bars) (B) Scatterplot of percent of sites under purifying selection vs percent of sites under positive selection in each protein. While overall the genome was under strong purifying selection, a substantial number of sites were identified as under positive selection in certain regions (eg, ORF3a, ORF10, N). (C) Sites detected to be under purifying selection (left panel) or positive selection (right panel) in spike protein. Gray areas indicate the NTD and RBD regions. (D) Sites under selection within the spike protein. Each amino acid residue is scaled from white (no selection) to blue (sites under purifying selection) and red (sites under positive selection) based on the probability of selection at the site. (E) Percentage of sites under purifying (top panel) and positive (bottom panel) selection per gene across eight time points (March 31 2020, June 30 2020, September 30, 2020, December 31, 2020, March 31, 2021, June 30, 2021, September 26, 2021) (F) Level of evidence (probability) of positive selection for (top-middel panels) spike sites with infectivity-associated HFV and (bottom panel) spike sites with escape-associated HFV.

When compared to other proteins within the SARS-CoV-2 genome, the spike protein and its RBD region were under some of the highest levels of purifying selection and lowest levels of positive selection observed, along with the M and Nsp7 proteins (Figure 2B, Figure S1A). However, spike sites 5, 142, and 950 were under some of the strongest diversifying forces in SARS-CoV-2 genome (Prob[alpha<beta] = 1, Figure 2C). The NTD was also enriched for sites under positive selection and depleted of sites under purifying selection when compared to the rest of the spike protein sequence (Figure 2B-C). While the function of the NTD is not entirely clear, neutralizing antibodies targeting a single supersite (shared residues across epitopes) in the NTD have been observed^39^. This supersite is enriched for diversification, with 9 out of the 34 (26%) of the residues under positive selection. Additionally, 11 sites within the RBD were under positive selection, including the six most frequently mutated sites (452, 478, 501, 484, 417, 477) (Figure 2C-D, Figure S1B). This indicated that despite having a globally conserved genome, evolutionary analysis at the single amino acid level revealed sites with significant diversity. A comprehensive list of all sites under selection is provided as a supplemental table (Table S1).

To better understand how these levels of selection have changed over time, we repeated this analysis across seven additional temporal windows (sequences collected up until March 31, 2020, June 30, 2020, September 30, 2020, December 31, 2020, March 31, 2021, and June 2021 respectively). We observed that most genes have undergone an intensification of purifying selection over time, with the exceptions of *ORF3a, ORF7a*, and *ORF10* (Figure 2E, Figure S1C). However, we observed strong positive selection at the sites of more transmissible variants, including 614^40,41^, 5^42^, 677^43^, 477^44^, and 501^45^ during the first year of the pandemic (2020) and more recently the delta-associated variant T478K^46^ (Figure 2F). Mutations that reduce neutralization by convalescent and therapeutic antibodies have also recently undergone intensified selection, including K417N/T^42,47,48^, E484K/Q^42,48-50^, and L452R/Q^42,50^ (Figure 2F-bottom panel). We also detected sites that were under diversifying forces earlier in the pandemic but no longer appear to be under selection, including the N439K mutation in the RBD (Figure S2).

### Impact of selective forces on SARS-CoV-2 infectivity and antibody neutralization

To further characterize how sites under selection within the RBD may relate to the pressures of the host environment, we first determined the impact of variants under selection on infectivity. We assessed the functional constraints of ACE2 binding using previously published deep mutational screens that measured the impact of mutations on affinity for ACE2 binding and protein folding stability, as estimated by RBD expression^51^. Substitutions at sites under purifying selection (83%) had the strongest negative impact on ACE2 binding affinity and protein stability (Figure 3A, Figure S3A). Conversely, sites under positive selection had less effect on binding or stability (Figure 3A). Since this screen was limited at estimating increased binding affinity due to global epistasis estimation, N501Y was the only HFV in the RBD that demonstrated a substantial increase ACE2 binding affinity in this analysis. However additional HFV have been reported to enhance binding affinity (N440K^52^, L452R^53,54^, S477N^44^, S477I^55^, T478K^46^, and S494P^56^), and these changes have been linked to increased infectivity and transmissibility^44,45,53,54^. We next looked to assess how sites under selection impacted antibody neutralization. We compared how the site-level estimates of selection related to the four major classes of SARS-CoV-2 neutralizing antibodies (Figure 3B). Each class varies in its ability to block ACE2 (class 1, 2) and its binding configuration to the RBD (class 1, 4: “up”-only; class 2, 3: “up” and “down”)^57^. While purifying selection applied to epitopes from all antibody classes, it had the strongest impact on epitopes from class 1 and class 4 antibodies (Figure 3C) and the lowest impact on epitopes from class 2 antibodies, which were reported to be associated with convalescent sera^50^. However, epitopes from each of these classes (with the exception of class 4) contained at least one site under strong positive selection which have been shown to impact therapeutic antibodies. Indeed, class 1 (e.g., casirivimab, etesevimab), class 2 (eg. bamlanivimab) and class 3 antibodies (e.g. C110) have been reported to be negatively impacted by mutations at the 417^58-61^, 484^58-62^, and 452^58,63-65^ sites respectively. The sites under positive selection significantly decreased antibody neutralization when compared to both sites under purifying (*p* = 0.009) and neutral selection (*p* = 0.006) (Figure 3D, S3B)^47-49,58,64-66^.

**FIGURE 3.**
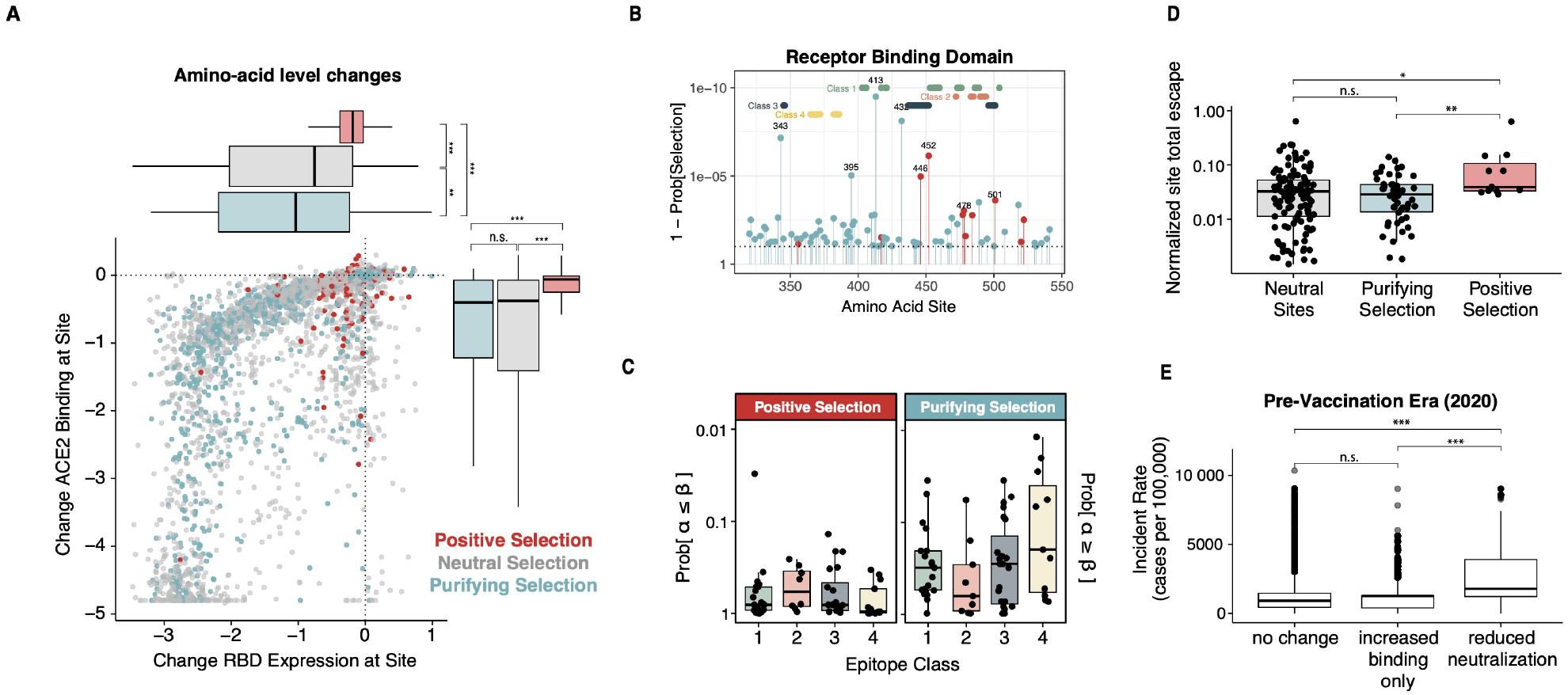
(A) For each possible amino acid substitution within the RBD, the effect of the mutation on RBD expression (x-axis) and ACE2 binding (y-axis). Each substitution is colored according to whether its associated site is under positive (red), purifying (blue), or neutral (grey) selection. (A) Sites detected to be under purifying selection (blue) or positive selection (red) within the RBD of the S protein. Overlayed are the sites associated with class 1 (green), 2 (orange), 3 (navy blue), and 4 (yellow) antibody epitopes. (B) Levels of positive (left) and purifying (right) selection of each antibody epitope class site (C) For each possible amino acid substitution within the RBD, the effect of the mutation on RBD expression (x-axis) and ACE2 binding (y-axis). Each substitution is colored according to whether its associated site is under positive (red), purifying (blue), or neutral (grey) selection. (D) Deep mutational scan normalized site total escape from convalescent plasma values for sites under positive (red), purifying (blue), or neutral (grey) selection. (E) Boxplot of the incident rates (cases per 100,000) for each sample in the pre-vaccination era (collected in 2020) with one of the RBD HFV in Table 1, grouped by whether that sample contains a mutation that reduces neutralization, increased ACE2 binding without any impact on neutralization, or no effect on neutralization and binding.

Finally, we analyzed the impact of all RBD HFVs at sites under positive selection and found that most (12/15) of these mutations have an impact on ACE2 binding affinity (e.g.: N501Y^67^), antibody neutralization (e.g.: E484K/Q), or both (L452R/Q^53,54,64^) (Table 1). Interestingly, we found that N501Y, E484K/Q and L452R/Q have all arisen independently across multiple lineages (N501Y in alpha, beta, gamma; E484K/Q in beta, gamma, iota, kappa; L452R/Q in delta, epsilon, kappa, lambda)^3^, indicating a common selective pressure acting on independent isolates. We also found that mutations at site 417 (K417N/beta; K417T/gamma) and 501 were physically linked in 95% of isolates (124,260/131,459). Because mutations at site 417 reduced antibody neutralization at a cost of ACE2 binding, and N501Y increased ACE2 affinity, the data suggested that some mutations can combine to alleviate the detrimental effect of individual variants.

Each of the currently characterized VUS contain at least one of the RBD HFVs that are associated with increased infectivity. Additionally, all except alpha also include a mutation that reduces antibody neutralization. The impact of the K417N/T and E484K on convalescent sera may have enabled the beta and gamma variants to spread in South Africa^68^ and Brazil^69^ despite high levels of seroprevalence in the late 2020 to early 2021 periods. To further investigate this, we annotated each isolate collected in 2020 with the incident rate (cases per 100,000) estimated from the geographic region and month from which the sample was collected, which we found to be a strong correlate with seroprevalence in the pre-vaccine era (Figure S4). We observed that the variants that reduced antibody neutralization were found in areas with significantly higher seroprevalence (Figure 3E). While most of the vaccines continue to demonstrate strong efficacy against the currently circulating variants, the beta and gamma variants have led to the discontinuation of both therapeutic antibodies (eg. bamlanivimab, etesevimab)^70^ and specific vaccines (e.g. ChAdOx1)^71^ due to loss of efficacy. Altogether, this suggests that the mutations associated with reduced neutralization have serious implications for public health.

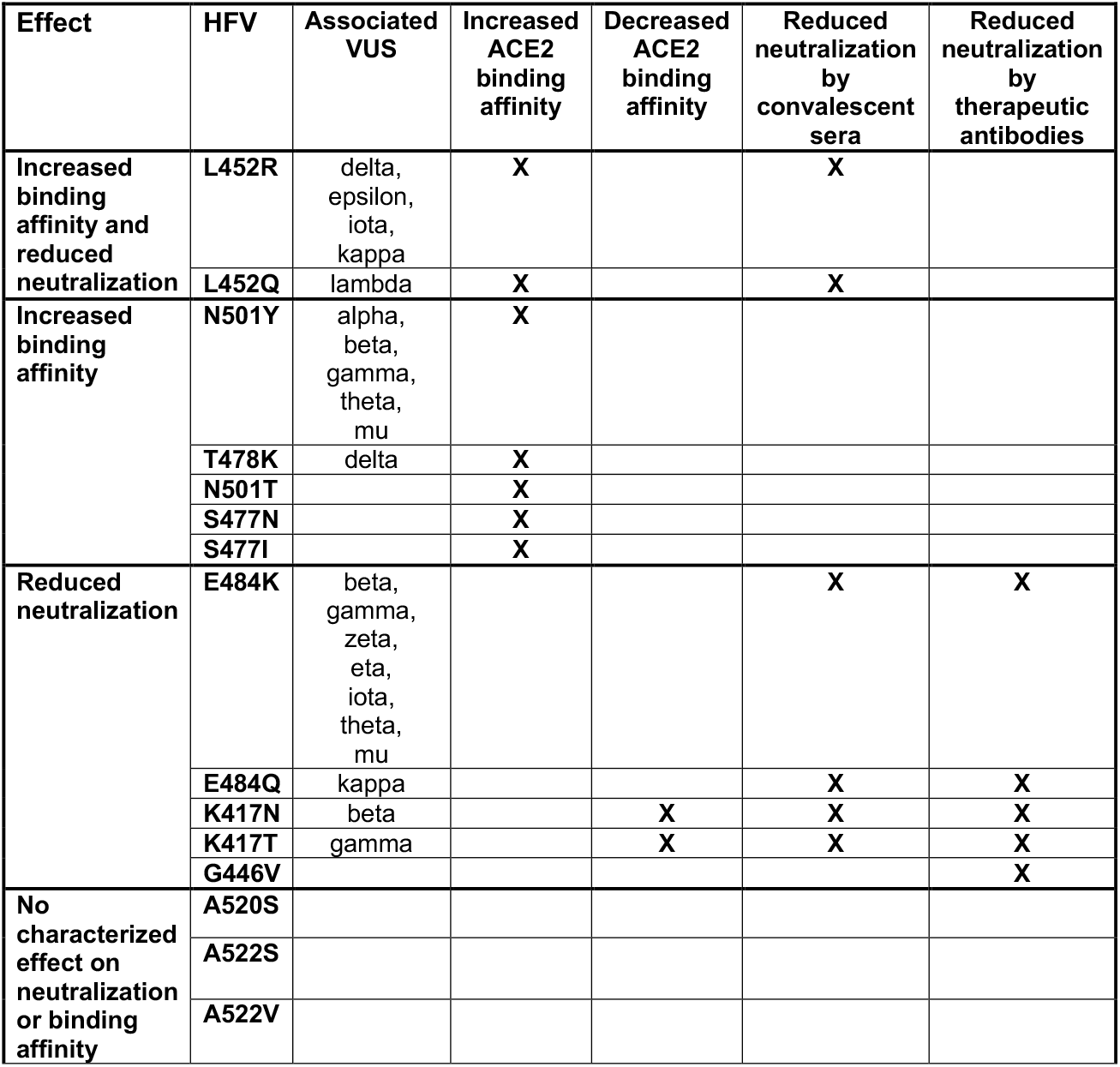

### Selective pressures within candidate variable T cell epitopes

In order to determine if SARS-CoV-2 mutations occur in regions targeted by the human immune system, we examined the distribution of amino acid mutations across 142 previously reported putative memory CD4^+^ T cell epitopes from SARS-CoV-2-naïve individuals^22^ and 637 previously reported candidate memory CD8^+^ T cell epitopes from both SARS-CoV-2-naïve individuals and recovered COVID patients^23^. For each peptide, we determined all amino acid mutations observed in SARS-CoV-2 genomes present in GISAID. While all peptides had each amino acid position mutated in at least one SARS-CoV-2 isolate, most epitopes had very little overlap with HFV (Figure 4A-B, Figure S5A-B). Interestingly, the exception to this was ORF3a, which had large overlap (>25% of sites) with HFV in 41% (13/32) of peptides.

**FIGURE 4.**
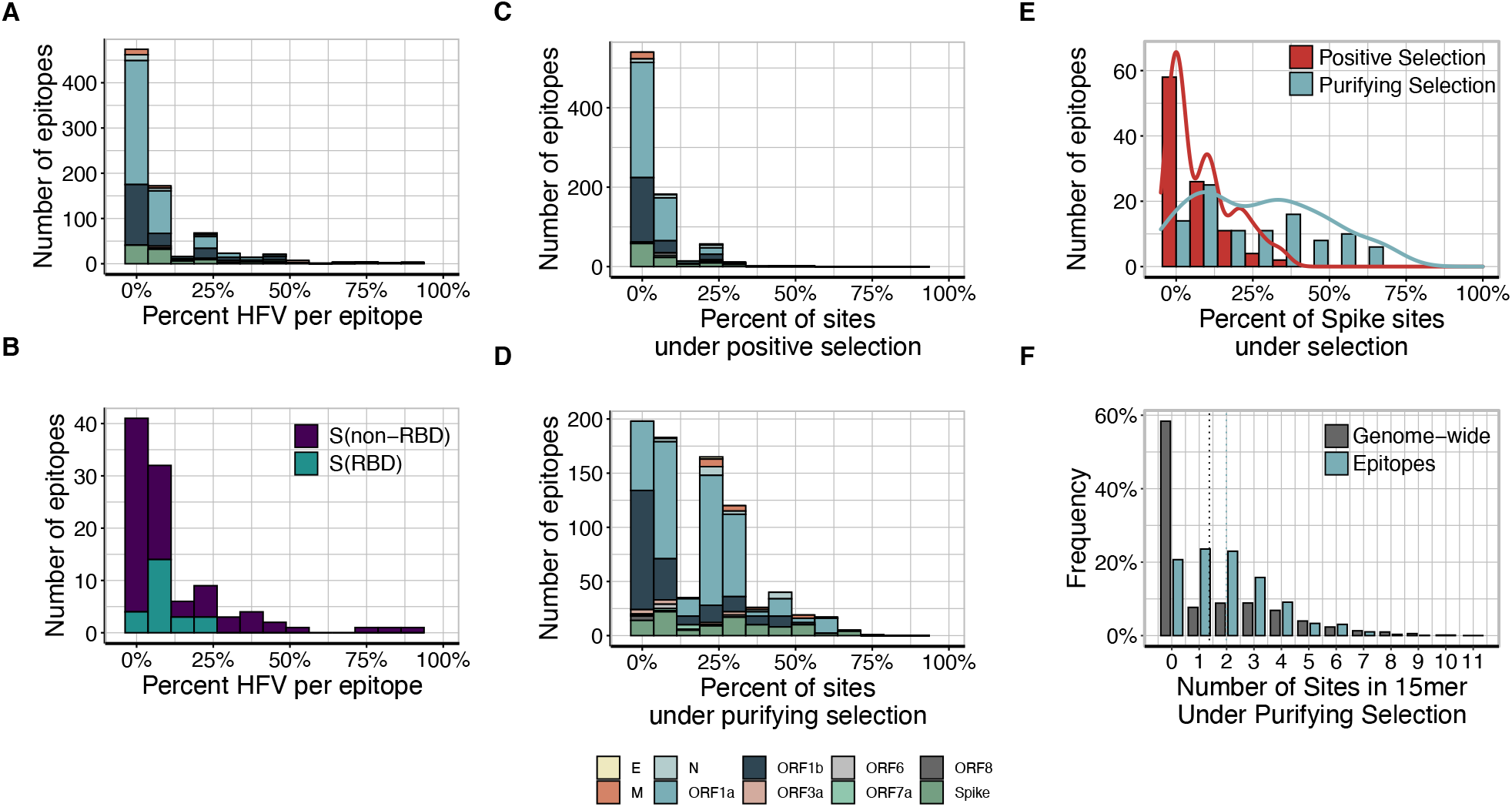
(A-B) Distribution of the number of sequence high frequency variants (found in 5000+ isolates) observed in (A) putative memory CD8+ T cell epitope sequences from recovered COVID patients, and (B) epitopes from RBD and non-RBD regions. (C-D) Number of putative memory CD8+ T cell epitope sequences from recovered COVID patients that contain sites under (C) positive and (D) purifying selection per gene (E) Distribution of the number of sites under selection in epitopes, broken down by positive and negative selection in spike peptides

To determine whether these putative T cell epitope sequences in SARS-CoV-2 strains were under selective pressure, we compared the number of sites under selection in the peptide sequences to background levels of selection. Overall, we found that T cell epitope sequences were significantly enriched for sites under purifying selection (*p* = 1.8×10^−67^, with a single epitope sequence on average containing 0-1 sites under positive selection and a third of their sites under purifying selection (Figure 4C-F). Notably, there were no significant differences in patterns of selection between either CD4 and CD8 epitopes nor between SARS-CoV-2-naïve and recovered COVID patients (Figure S5C-G). In addition to the absence of diversifying mutations, the spike, *ORF1a*, and *N* epitope sequences were significantly enriched for sites under purifying selection compared to the full gene (spike *p* = 1.9×10^−16^, ORF1a *p* = 4.9×10^−81^, N *p* = 1×10^−8^) (Figure S6). Overall, the data indicates that T cell epitope sequences are under strong purifying selective pressures.

## DISCUSSION

A significant finding of the present study is the strong evidence that SARS-CoV-2 antigen and epitope conservation is the rule, not the exception, in this highly successful human pathogen. The SARS-CoV-2 RBD is both the region mediating viral entry into the cell and the main target of potent neutralizing anti-spike antibodies in COVID-19 patient sera or plasma samples^72,73^. Therefore, most of the selective pressure imposed by the immune system of the host should target this domain. Close examination of RBD conservation indicated that while the RBD as a whole was under extensive purifying selection, consistent with the functional constraints imposed on this region, there were individual sites under strong levels of diversification. We found that these sites were associated with increased ACE2 binding and infectivity, as well as reduced neutralization by convalescent and therapeutic antibodies.

The HFV at position 417 of the RBD (K417N/T) presented a particularly interesting case. While these mutations reduce neutralization of therapeutic and convalescent antibodies^47,58^, this occurs at the fitness cost of reduced ACE2 binding affinity. We found that mutations at this site have not been successful in spreading without the addition of the compensatory mutation N501Y. Experimental studies have confirmed that while K417N alone results in a significant drop in ACE2 binding affinity, its combination with N501Y resulted in 3-fold stronger binding than wild-type (although still 2-fold less compared to N501Y alone)^74^. More recently, K417N has also been found in two sub-lineages of delta (AY.1, AY.2). While these variants do not include N501Y, they do include a different compensatory mutation that increases ACE2 binding affinity, L452R. Neutralizing monoclonal antibodies have also been isolated that target non-RBD epitopes (Chi X., et al.,2020; Liu et al., 2020). For example, antibodies 4A8^75^ and 1-68^18^ have been reported to neutralize SARS-CoV-2 by targeting the NTD. However, these NTD-directed neutralizing antibodies have been found to target a single supersite^39^, which was characterized by elevated levels of positive selection. A number of HFV also overlap with the NTD supersite, including the 144 deletion found in the alpha variant, the 242-244 deletions in the beta variant, and multiple NTD mutations found in the delta variant. These variants in turn have demonstrated reduced sensitivity to anti-NTD neutralizing antibodies^76^.

Another significant finding of the present study is the identification of a substantial number of sites under both positive and purifying selection outside of the spike protein. We found elevated and intensified levels of diversification in the *N, ORF3a, ORF7a, ORF8, and ORF10* genes. This can be explained by the roles these genes have been described to play in interacting with the immune system to block the innate immune response. Additionally, the N R203M mutation associated with the delta variant was described to significantly increase mRNA production and delivery in to host cells and thus explain the increased transmissibility of this successful variant^77^. In our analysis, R203M mutation site was identified as being under selection, demonstrating that identification of sites under selection and subsequent phenotypic characterization of mutations occurring at these sites can help better understand the keys of SARS-CoV-2 success. Altogether, this suggests that the pressures outside of the spike are also important for fitness and warrant closer examination.

We also examined experimentally verified peptide epitopes derived from unexposed donors^22,23^ and recovered COVID patients^23^ for sequence variants in SARS-CoV-2 and found that the vast majority were under strong conservation and did not accumulate sites under positive selection. In uninfected donors, SARS-CoV-2-reactive T cells can exhibit different patterns of immunodominance and frequently target *ORF1a* coding sequence, *nsp7*^10^. In the present study, we showed that *nsp7* is hyper-conserved and has no sites actively under positive selection. This observation is consistent with the high level of identity between orthologous sequences of *nsp7* across multiple coronaviruses^78^ and helps explain why *nsp7*-specific T cells are detected and often dominant in unexposed donors as they may derive from exposure to other coronaviruses. By contrast, previous studies report that *nsp7*-specific T cells usually represent a minor population in individuals who recovered from COVID-19^10^. These patients often have T cells preferentially reacting to structural proteins, including N protein-derived peptide pools^10^.

Since September 1^st^ 2021, the vast majority (>99.6%) of collected SARS-CoV-2 sequences have been of the delta lineage. The majority of existing HFV mutations have been found in a substantial number (>500) of delta isolates, including K417N and E484K, which we observed to be under convergent evolution. Additionally, 30 sites associated with delta lineage defining mutations were identified as under positive selection by our analysis, including 17 sites in the spike and 8 sites in the N and *ORF3a* genes (4 sites each).

In conclusion, our findings indicated that SARS-CoV-2 genetic selection is driven by a deterministic process imposed by natural within-host purifying selection, leading to a dominant lineage mode of evolution. However, our results also showed that this pathogen is capable of genetic plasticity dictated by environmental changes. The necessity to adapt leads to selection and the contribution of dominant genotypes determined by the host.

## METHODS

### Sequence analysis, filtering, and alignment

The approximately 4.7 million complete genome sequences deposited in GISAID^1,2^ as of September 26, 2021 were downloaded along with their variant calls and accompanying metadata. GISAID genomes were filtered to remove poor quality sequences using the following criteria: complete metadata, less than 2% ambiguous DNA sequence (N) or protein (X) with no runs of 4 or more Ns in the genome or Xs in the proteins, all genes (ORFs) in the genome are matched by a BLAST search and were not truncated by premature stop codons. The premature stop codon constraint was relaxed for ORF8, which has a stop codon at amino acid position 27 in the widespread ^34^1^35^1. To obtain full length coding genomes, coding sequences of each open reading frame were aligned with ^79^11^79^1 The genome sequences that passed QC were also filtered to remove accessions that are not represented in the September 26^80^1 downloaded from GISAID, 427,773 sequences passed all of the QC filters and were also in the Audacity tree. To obtain a more computationally tractable data set, a subset of 20,000 sequences was selected using random down-sampling. To ensure that the novel variants were represented, a minimum of 100 variant sequences, where available, were preserved in the subsample. The Audacity phylogenetic tree was pruned to contain only the accessions that passed QC, then further pruned to match the 20,000-sequence subsample. The resulting phylogenetic tree was used as the guide tree for selection and further phylogenetic analysis. The full set of 427,773 QC-passed sequences was also used to generate quarterly time-point data sets. For each time-point (2020-03-31, 2020-06-30, 2020-09-30, 2020-12-31, 2021-03-31, 2021-06-30, 2021-09-26 the sample collection date was used to generate a subset of sequences that were collected up to that date. For the time points with > 20,000 QC-passed sequences, down-sampling was performed as described above.

### Phylogenetics

The downsampled subset of 20,000 genomes and the corresponding portion of the Audacity tree were used to perform a further, phylogeny-guided subsample of 1000 genomes with Treemmer^81^ to reduce the sample size while preserving sequence diversity. Where available, at least 10 representatives of the variant lineages were protected from down-sampling to ensure they were represented in the final tree. A whole-genome maximum likelihood tree of the 1000 sequences was constructed using iqtree2^82^ with the GTR+F+R3 model. The phylogenetic tree was outgroup rooted using a related bat betacoronvirus genome (RATG13; GISAID accession EPI_ISL_402131) to infer the evolutionary branching order of SARS-CoV-2 lineages. Phylogenetic trees were annotated with geographic and variant metadata using the ETE3 Python toolkit^83^. Images of annotated phylogenetic trees were rendered using the Python API to <u>iTOL</u>, the interactive Tree of Life^84^.

### Natural selection analysis

The selection test was performed using Fast Unconstrained Bayesian AppRoximation (FUBAR) method^30^ from the hypothesis testing using phylogenies (HyPhy) Suite (https://stevenweaver.github.io/hyphy-site). The analysis was conducted for each single gene. The prepared alignment and corresponding phylogenies described above were sent to FUBAR to infer nonsynonymous (dN) and synonymous substitution (dS) rates at a per-site basis and test whether dN was significantly different from dS. Sites with probability values > 0.9 were considered to be under non-neutral selection.

To assess whether the genome subsets were representative of the larger data set and that any results will be robust and reproducible, his analysis was repeated across five different replicates of the above-described downsampling procedure. The consistency of the results was assessed using Pearson correlation of the log(1 -probabilities) of positive selection across the full SARS-CoV-2 genome. There was a high degree of concordance among replicates (Figure S7). This suggested that these downsampled subsets were sufficiently representative of the larger dataset and could be used to further investigate the evolution of the SARS-CoV-2 genome.

### Spike Protein Visualization

Using the output of variant calling scripts, we identified how many viruses were sequenced that contained a mutation from the wild type sequence at each position. These frequencies were used to annotate the B-factor of PDB file 6VSB^85^ with PDB ID 6M0J^86^ with PDB ID 6M0J^86^ superimposed on one protomer of 6VSB in order to provide resolution on residues in VUS which lacked resolution in 6VSB; The resulting chimeric structure was used to color each atom using PyMOL’s *spectrum* command.

### Analysis of deep mutation scan data

Data from deep mutational scans was downloaded from https://github.com/jbloomlab/SARS-CoV-2-RBD_DMS (RBD binding, expression constraints) and https://github.com/jbloomlab/SARS2_RBD_Ab_escape_maps (escape maps). The escape value was taken to be the average of “normalized_site_total_escape” for site-level and average of “mut_escape” for mutation-level across all convalescent plasma samples.

### Seroprevalence analysis

CDC seroprevalence estimates was downloaded from https://covid.cdc.gov/covid-data-tracker/#serology-surveillance. The John Hopkins daily incident rate estimates were downloaded from https://github.com/CSSEGISandData/COVID-19. For each US state, the relationship between seroprevalence estimates and incident rates was evaluated using Pearson’s correlation coefficient. Since these were highly correlated (R=0.8), the incident rate was used as a proxy for seroprevalence. For each sample in GISAID, the incident rate at the month (Collection Date) and highest-resolution place (county when available, otherwise state/country) was annotated. The incident rates were used to compare trends where different sets of mutations are spreading. This analysis was restricted to the pre-vaccination era of sequences, as defined by the samples collected between June 2020 (when incident rates became reliable) and December 31, 2020 (when vaccinations began to ramp up in the United States).

### Analysis of CD4+ and CD8 + T cell epitope sequences

The list of 142 previously published memory CD4^+^ T cell epitope sequences and genome coordinates was downloaded from Supplementary Table 1 of Mateus et al., 2020^22^. The list of 637 previously published CD8^+^ T cell epitope sequences were downloaded from Supplementary Tables 4 and 7 of Saini et al., 2021^23^ and each peptide sequence was blasted against the reference sequence (accession: MN908947.3) to determine genome coordinates.

For each gene, we counted up the total numbers of unique sites that were covered by epitope sequences and under positive/purifying selection. Significance was assessed using a Chi-square test comparing the number of sites under selection within sites targeted by candidate epitopes to the sites within the same gene/protein not targeted by candidate epitopes. The number of sites under positive/purifying selection were computed for all 15-mers genome-wide and compared to epitope sequences using the one-sided Wilcoxon rank-sum test for each gene. The number of sites under positive/purifying selection were also computed for all 15-mers genome-wide and compared to epitope sequences using the one-sided Wilcoxon rank-sum test for each gene.

## SUPPLEMENTAL FIGURE LEGENDS

**SUPPLEMENTAL FIGURE S1.**
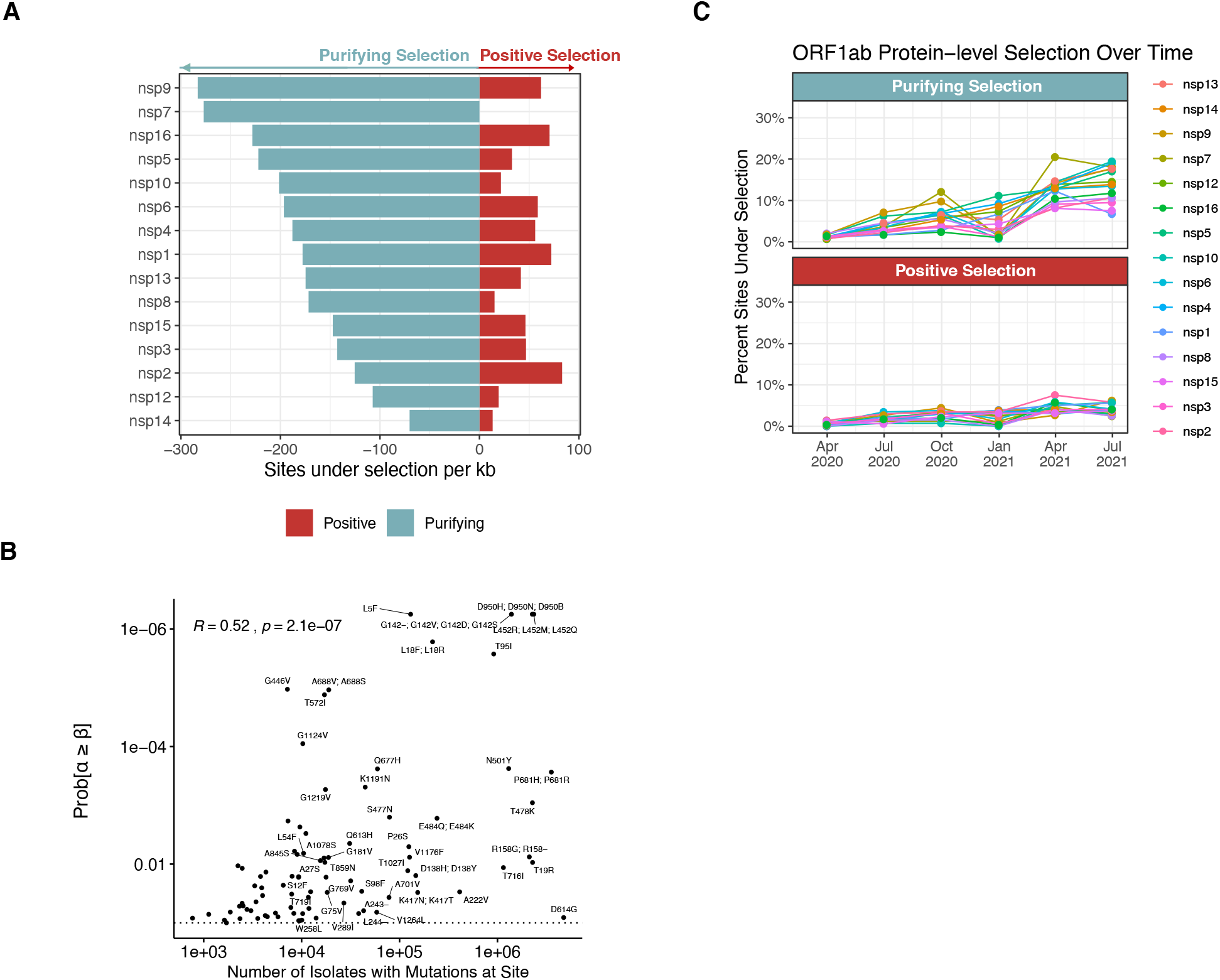
**Related to Figure 2**. (A) Protein length normalized counts (counts per kb) of sites under significant purifying selection (light blue bars) and positive selection (red bars) for the protein products of ORF1ab gene. (B) Scatterplot of the probability of positive selection at a site versus the number of isolates at that site with mutations. (C) Percentage of sites under purifying (top panel) and positive (bottom panel) selection per ORF1ab protein product across seven time points (March 31 2020, June 30 2020, September 30, 2020, December 31, 2020, March 31, 2021, June 30, 2021, September 26, 2021)

**SUPPLEMENTAL FIGURE S2.**
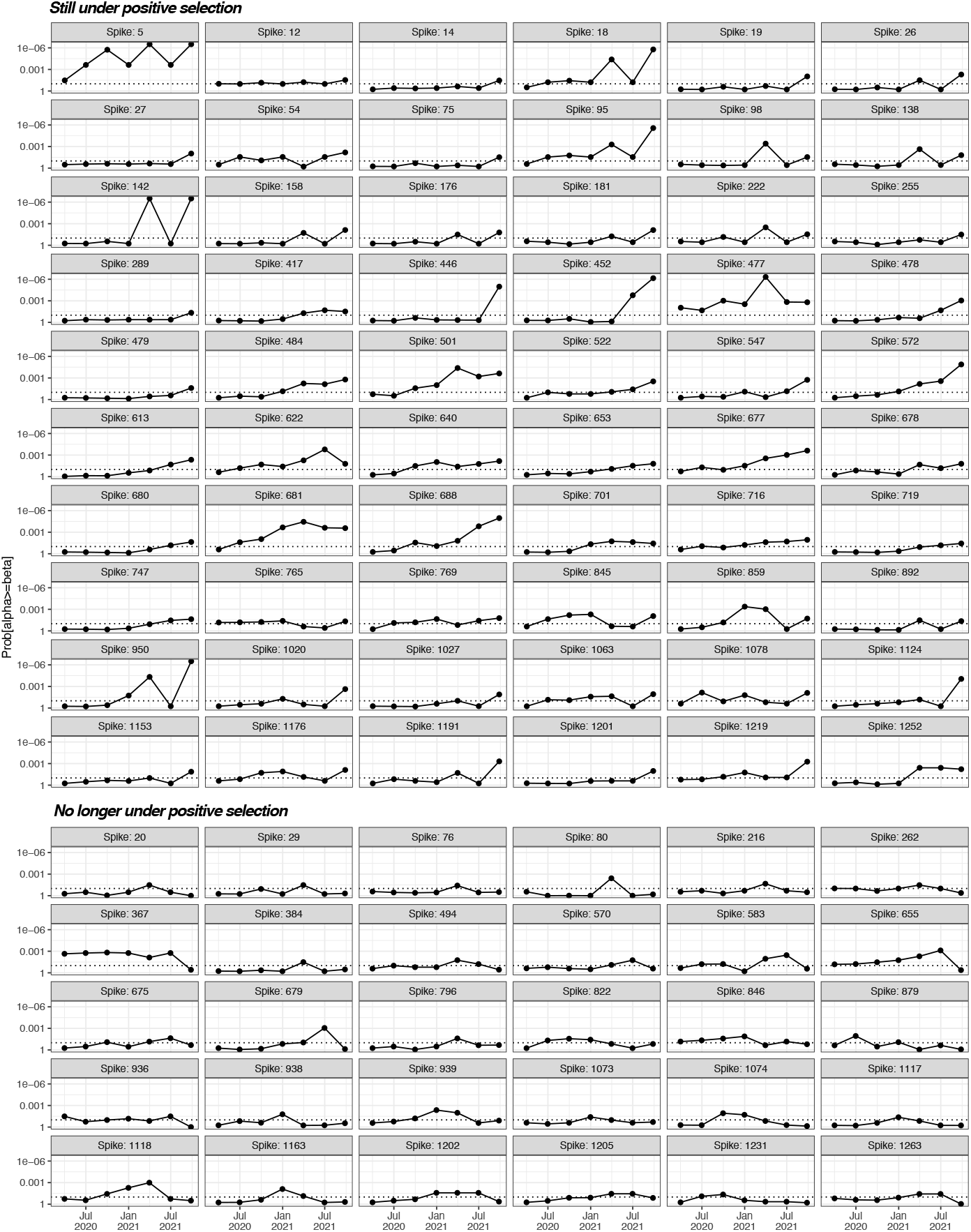
**Related to Figure 2**. Probability of selection for the top 90 spike sites that have undergone positive selection, grouped by whether they were under selection at the September 26 analysis date (top) or no longer under selection (bottom)

**SUPPLEMENTAL FIGURE S3.**
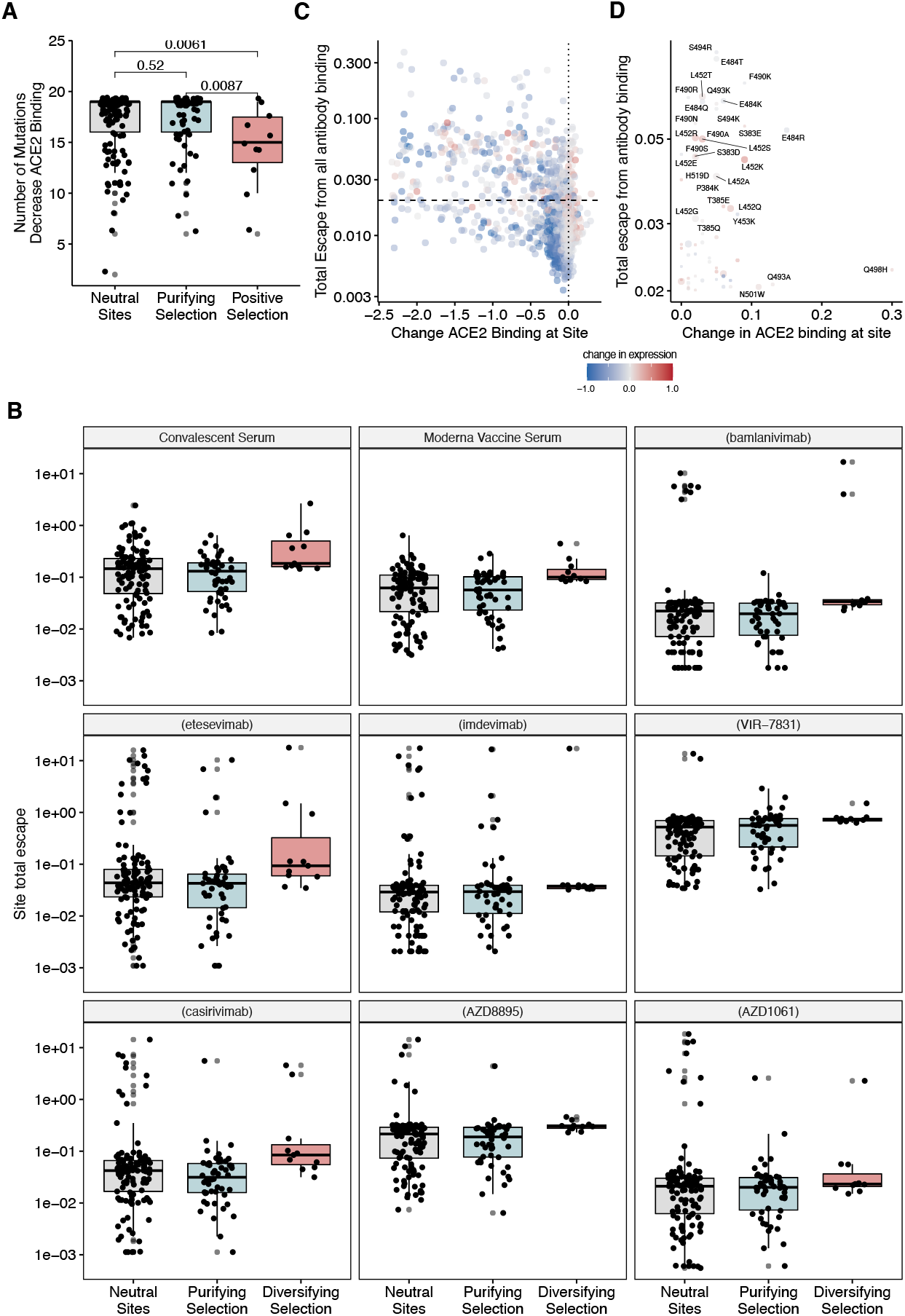
**Related to Figure 3**. (A) Boxplot of the number of possible amino acid substitutions that decrease ACE2 binding for each position in the RBD, grouped by whether the site is under positive, purifying or neutral selection. (B) Deep mutational scan site total escape values from convalescent plasma, antibodies from Moderna vaccinated patients, and therapeutic antibodies for sites under positive (red), purifying (blue), or neutral (grey) selection. (C-D) For each possible amino acid substitution within the RBD, scatterplot of the effect on ACE2 binding, effect on RBD expression, and average escape from convalescent antibodies.

**SUPPLEMENTAL FIGURE S4.**
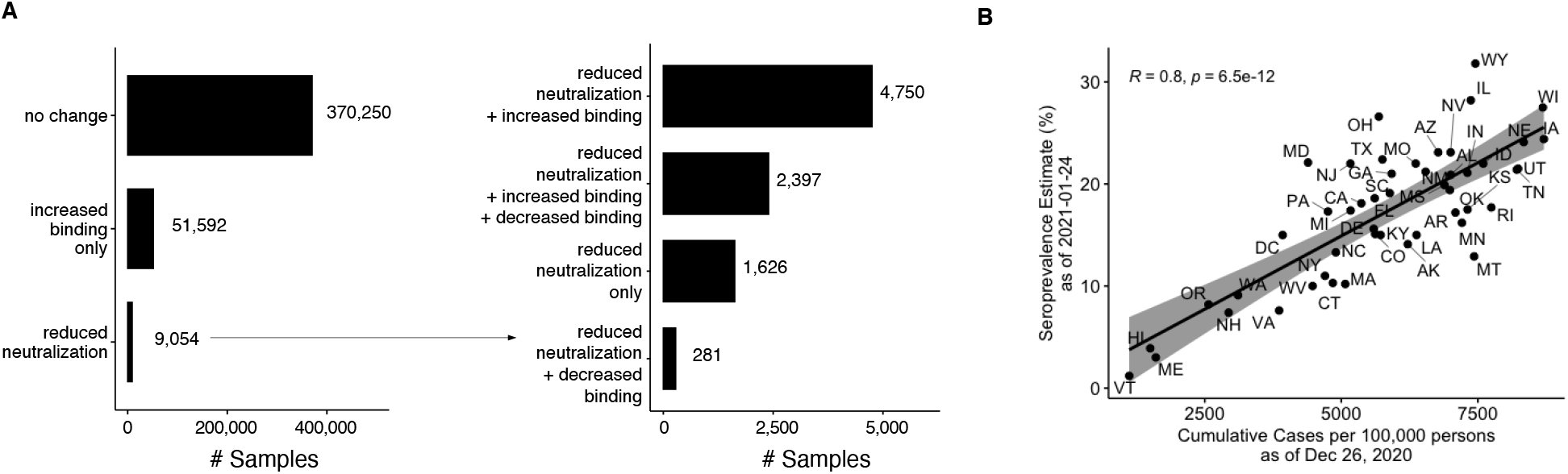
**Related to Figure 4**. (A) Number of isolates with any combinations of RBD mutations which confer increased ACE2 binding, escape from convalescent antibodies, loss of ACE2 binding (loss), or neither (no change). (B) For each US state, correlation of the CDC seroprevalence estimates from January 24 with the incident rate (cases per 100,000) corresponding to three weeks prior (December 26)

**SUPPLEMENTAL FIGURE S5.**
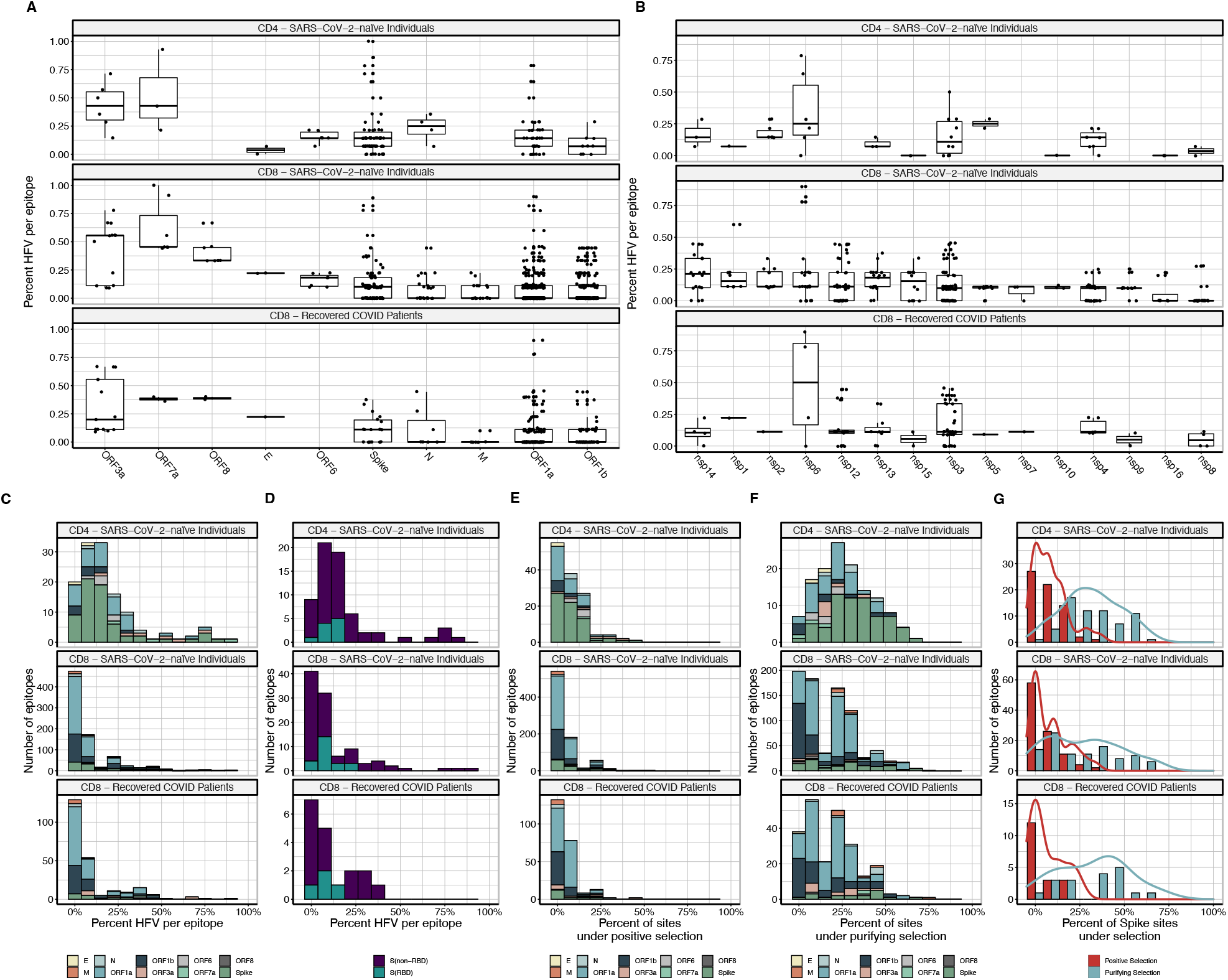
**Related to Figure 4**. (A-B) Percent of epitopes with high frequency variants (HFV), broken down by (A) gene and (B) ORF1ab protein product (C-D) Distribution of the number of sequence high frequency variants (found in 1000+ isolates) observed in (A) putative memory CD8+ T cell epitope sequences, and (B) epitopes from RBD and non-RBD regions. (E-F) Number of putative memory CD8+ T cell epitope sequences that contain sites under (E) positive and (F) purifying selection per gene (G) Distribution of the number of sites under selection in epitopes, broken down by positive and negative selection in spike peptides (top panel) putative CD4 T cell epitopes from SARS-CoV-2 naïve individuals (middle panel) putative CD8 T cell epitopes from SARS-CoV-2 naïve individuals (bottom panel) putative CD8 T cell epitopes recovered COVID patients

**SUPPLEMENTAL FIGURE S6.**
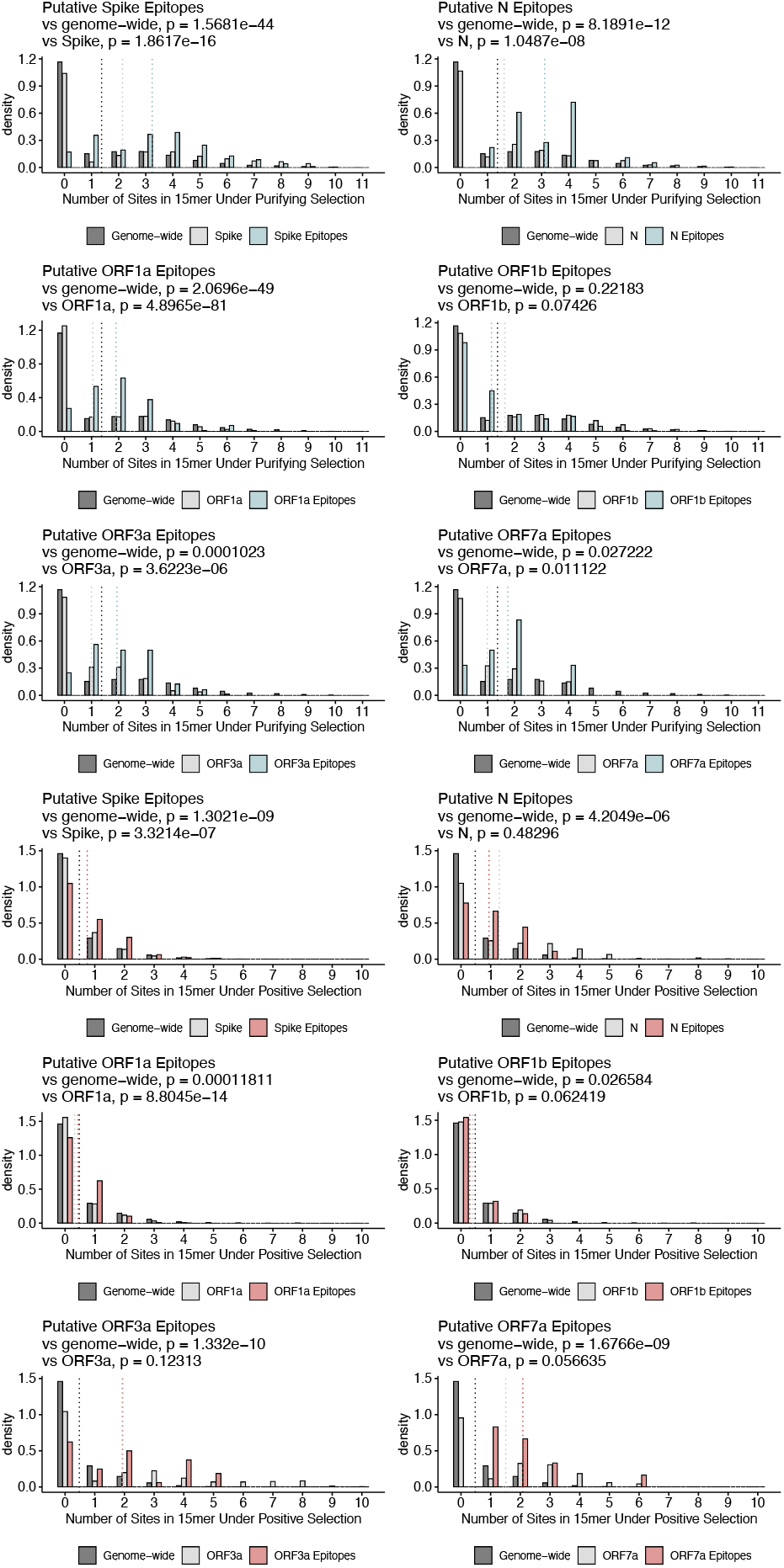
**Related to Figure 4**. Distribution of the number of sites under purifying and positive selection in spike, N, ORFa1, ORF1b, ORF3a, and ORF7a peptides and genome-wide 15-mers. Significance was assessed using a one-sided Wilcoxon rank-sum test.

**SUPPLEMENTAL FIGURE S7.**
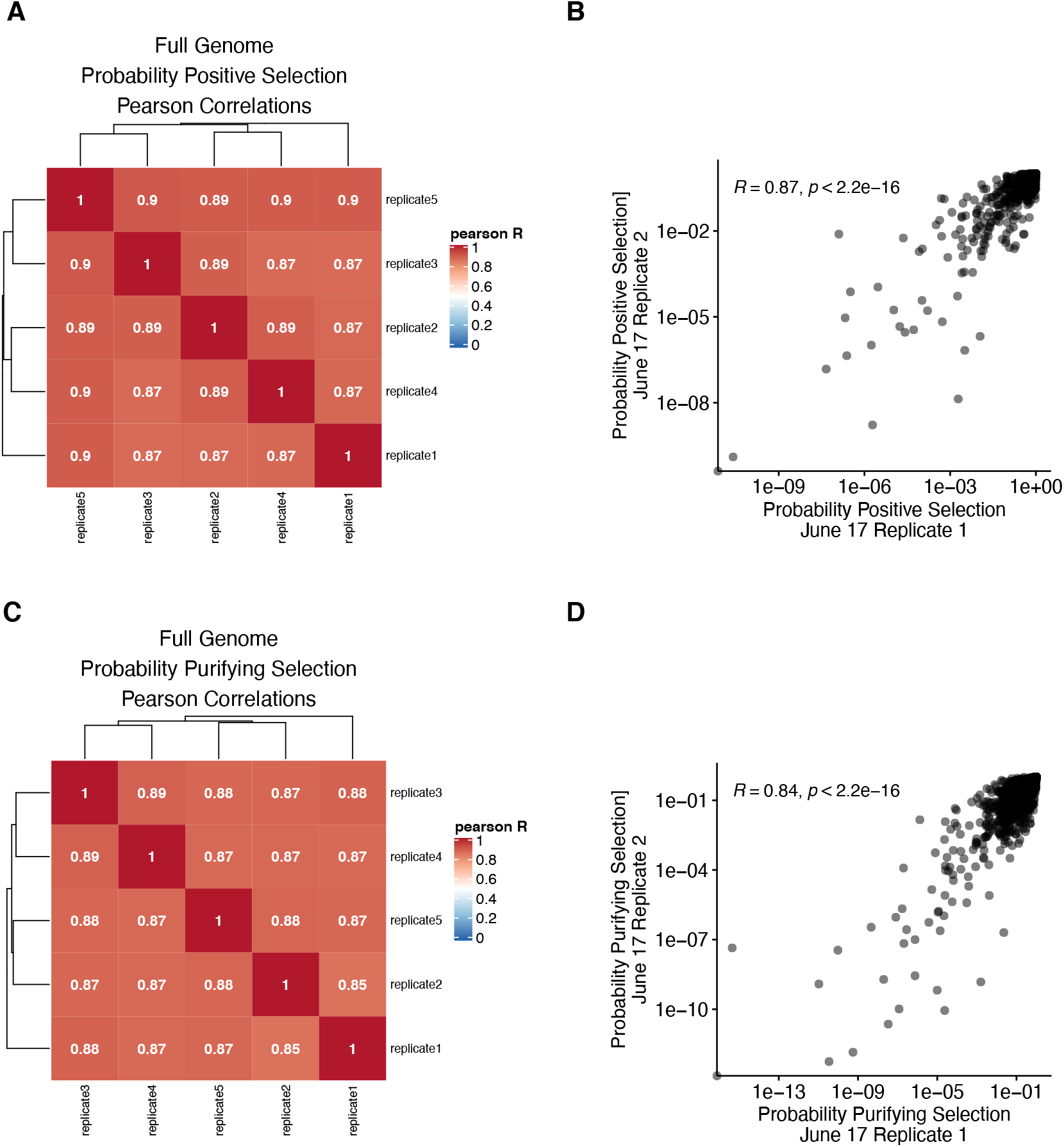
**Related to Figure 2**. (A) Pearson correlation coefficients of the log probabilities of positive selection across five different down-sampled replicates (B) Scatterplot of the log probabilities of positive selection between two down-sampled replicates (C) Pearson correlation coefficients of the log probabilities of purifying selection across five different down-sampled replicates (D) Scatterplot of the log probabilities of purifying selection between two down-sampled replicates

